# Identifying a logical specification and a program for an LLM-based generator of lead molecules

**DOI:** 10.1101/2025.02.14.634875

**Authors:** Ashwin Srinivasan, Tirtharaj Dash, A Baskar, Sanjay Kumar Dey, Mainak Banerjee

## Abstract

Our interest is in the generation of “lead” molecules in early-stage drug design. Leads are small molecules (ligands) that can bind to a part of pre-specified target and also satisfy multiple physico-chemical constraints. We propose using techniques developed in Inductive Logic Programming (ILP) to identify a logical specification of feasible molecules; and then using this specification to construct a program that uses a large language model (LLM) to generate new molecules. We ensure the program constructed is correct, in the sense that every molecule generated by the program is feasible according the specification. Our focus is on contributing to on-going drug-discovery research on novel inhibitors for Dopamine *β*-hydroxylase (DBH), an enzyme that plays a pivotal role in several diseases related to the brain and the heart. We find molecules comparable in affinity to the latest generation drugs currently in clinical trials, and chemical assessment of synthesisablity of the molecules generated. For completeness, we also provide results obtained on the classic benchmark datasets used in recent work reported in [1].

## 1 Introduction

The use of AI for scientific discovery has been with us at least since DENDRAL [2]. By this we mean models that can provide explanations for, and make predictions about, observational data. The broad goal has been to accelerate the interplay between conjecture and criticism that captures the essence of scientific reasoning (see Fig. 1(a)). Our interest is in the use of machine-learning (ML) for the identification of new drugs. The development of new drugs, with its well-documented scientific complexity, economic costs, and social need, presents an ideal case for the need for a science accelerator. The tasks involved are shown in Fig. 1(b).

**Figure 1.**
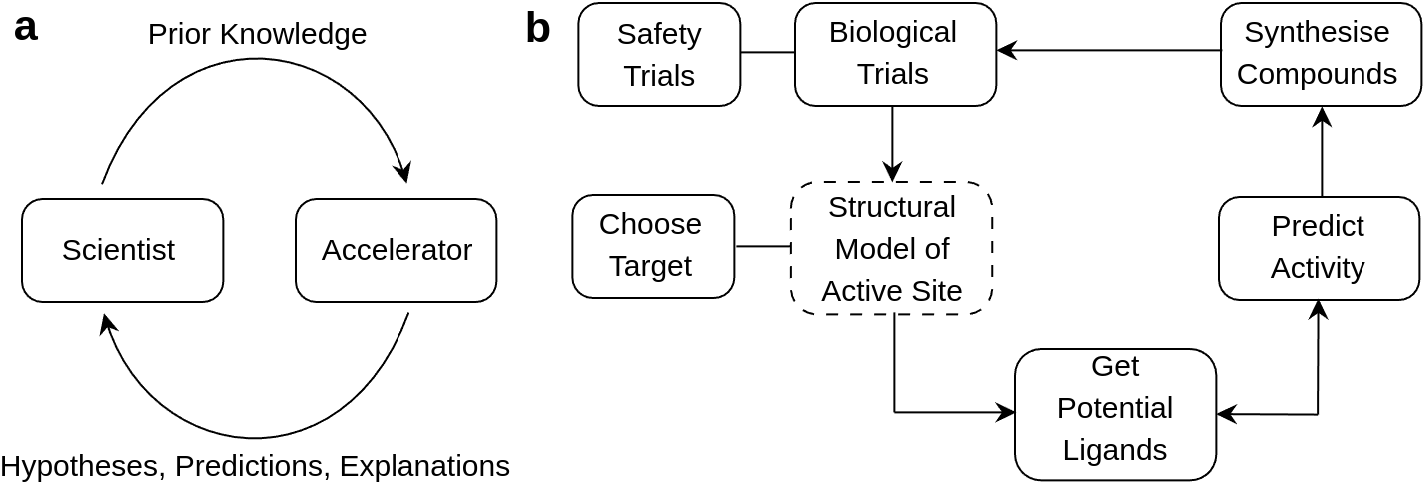
(a) An automated scientific assistant intended to accelerate the process scientific discovery (not shown is that both human and machine can access external data and information sources); (b) Some of the main tasks in drug-discovery. A target is usually a protein, and the active site refers to some part of the protein that acts as a “lock”. A ligand is usually a small molecule that acts as a “key” that fits into the lock. Ligands that satisfy additional chemical constraints are called leads. Leads that satisfy additional biological and safety constraints may result in drugs. The tasks of choosing a target and getting a ligand are inherently generative and that of predicting activity is discriminatory.

Perhaps the most complete demonstration of a science-accelerator using ML with potential applications to drug-discovery, can be found in the development of a Robot Scientist [3]. An important feature of this work is the provision and use, by the ML approach, of prior scientific knowledge. ML used to construct an updated hypothesis as explanation, along with testable predictions. The re-use of formal representations of prior knowledge in the Robot Scientist employs symbolic ML, and relatively narrow range of biochemical knowledge. However with recent advances in ML, we have access–for example through the use of large language models, or LLMs–to vast stores of (approximate) knowledge on a variety of areas relevant to several parts of the drug-design cycle. Thus it is possible to envisage constructing accelerators that are able to generate hypotheses about potential drugs that not only account for structural constraints (as is done in the Robot Scientist), but also constraints on synthesis and prior results from biological testing.

More recently, it was shown in [1] that an LLM-based ligand generator could be progressively aligned to generating molecules that high estimated affinity to a target. This was done using a sequential approach to identifying the appropriate conditional distribution for the LLM by an iterative process inspired by MIMIC [4], a model-driven estimation of distribution algorithm (EDA: [5]). This focussed the distribution used by the LLM on molecules with progressively higher affinity estimates. The procedure was tested on two well-known target molecules, for which substantial chemical knowledge is known. Computational chemists were able to confirm that the LLM-generated molecules were novel, and did have the kinds of structural features needed for inhibiting the targets. This showed at least 3 things: (1) LLMs appear to already have the kinds of biochemical knowledge needed to bind to the target; and (2) It is possible to use a MIMIC-style EDA to provide the contextual updates required for the LLM to narrow down quickly on the relevant biochemical knowledge; and (3) LLMs are usually able to generate valid “molecular sentences” (in this case, SMILES strings). There are however some limitations of the work in [1]. First, sequential sampling using the LLM is based on progressive change of a single numerical property of the molecule. namely: estimated binding affinity of the molecule to the target. While high binding affinity is desirable, it is not the only pre-requisite to be satisfied by a lead molecule. In practice, we need to generate molecules subject a variety of chemical, biological and economic constraints. Secondly, it is assumed that we have access to a sequence of thresholds on affinity that are used to progressively alter prompt, and hence the context for the LLM. In effect, we require prior knowledge of an optimisation schedule. This may be possible with a single constraint like binding affinity. However as the number of constraints increase, the requirement becomes increasingly difficult to satisfy. Finally, the experiments were restricted to well-studied targets for which there are already known leads. In this paper, we address each of these shortcomings.

Recent advances in AI, especially large language models (LLMs) have significantly impacted drug discovery by facilitating molecular design and lead identification [6, 7]. The trending models for AI-assisted drug design are primarily based on deep neural networks. Until, a few years ago, the classic generative model of choice used to be variational auto-encoder (VAE) [8] and generative adversarial networks (see [9]). In addition, there are some work that focus on generating molecules using VAE while also integrating domain-constraints (see [10], [11]). In [11], the generative model is a combination of two VAE-based generators and a discriminator (Bot-GNN [12]). In that work, it was shown that domain knowledge played a crucial role in assisting lead discovery by embedding expert-defined rules within GNN via bottom-clauses constructed by mode-directed inverse entailment in ILP [13], ensuring the generated molecules adhere to chemical synthesis feasibility and biological relevance. Although, classical VAE based generative models are still being used, the focus has largely shifted to large language models in the last few years. For instance, the precursor to the current work uses LLMs to generate novel molecules using domain-knowledge and constraints encoded as a logical formula. The logical feedback mechanism allows the LLM to iteratively generate and refine molecular candidates based on chemical and biological constraints, improving their viability as potential leads [1] for target proteins. In a similar direction, TamGen [14] uses a GPT-like chemical language model to enable target-specific molecule generation and refinement, improving molecular quality and viability. Their focus was on generating novel inhibitors for Tuberculosis ClpP protease. In [15], the authors show that protein structure can assist generating better inhibitors. Even even though there was no experimentally determined structure known for a drug target, the structure generated by AlphaFold [16] can still be useful.

The main contributions of this paper are: (1) We propose a systematic methodology that integrates the concept of Inductive Logic Programming (ILP) with Large Language Models (LLMs) for molecular generation; (2) We provide formal specification of our approach and show that the program constructed is correct, that is, every molecule that the LLM generates are valid and adhere to the required physico-chemical constraints; (3) We demonstrate its application of this approach to discovering novel inhibitors for three different target proteins: two kinase inhibitors (Janus Kinase 2, Dopamine Receptor D2) and Dopamine *β*-Hydroxylase (DBH). Our proposed approach explores a multi-dimensional search space encompassing various factors relevant to molecule generation, including physico-chemical properties, unlike previous methods that rely solely on numerical constraints such as affinity thresholds [1]. Furthermore, we use two domain-experts to critically assess the generated molecules and provide their views on their synthesisability.

The rest of the paper is organised as follows. In section 2, we provide conceptual description of constructing a logical program for generating molecules with more than one factor. In section 3, we provide emprical evaluation of our methodology when applied to three proteins and assessments by two domain experts. We conclude the paper in section 4.

## 2 Multi-Factor Molecule Generation

It is evident from Fig. 1(b) that even a single iteration of the cycle of identifying, synthesising and testing a potential drug will require the molecule to pass through several stages. The structural constraint of target-specific binding is necessary but not sufficient. Additionally, there are target-agnostic physico-chemical constraints (for example, on molecular weight, hydrophobicity *etc*.), synthesis constraints (the number of steps required for synthesis, yield), cost of manufacture, mutagenicity in chemical and biological tests and so on. Ideally, we would like any sample of new molecules to be drawn from a distribution that is likely to satisfy all such constraints. In practice, we would settle for a sample that contains molecules that are a good compromise on the criteria. The mathematical problem is therefore one of finding satisfactory solutions through a process of optimisation.

In this paper, we construct a program for molecule-generation two stages, motivated by the classic software-engineer’s approach of first identifying a specification, and then using the specification to obtain an implementation. Unlike traditional software engineering, however, our specification will be constructed automatically, using techniques from Inductive Logic Programming (ILP: [17]). A randomised search is used to arrive at a logical definition for the set of feasible molecules. The logical definition then used to obtain an LLM-based implementation that includes a form of automatic prompt restructuring by exploiting the utility of in-context learning for LLMs [18].

### 2.1 Stage 1: Data to Logical Specification

The first stage is concerned with identifying an appropriate logical description for feasible molecules. In this paper, we will assume the set of “feasible molecules” is specified by a conjunction of constraints. For example:

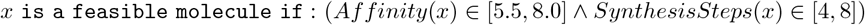

On the face of it, any method for learning axis-parallel (hyper-)rectangles would appear to suffice to identify conjuncts of the kind shown above.^1^ The difficulty arises not in principle, but in what is practice. First, for any interesting drug-design problem, the number of feasible molecules are usually very few in number, and we want methods that can work even with 1 molecule. Secondly, we want the approach to go beyond the molecules that are known. Thus, techniques that construct descriptions that are just based on the data available are unlikely to yield interesting results (this applies also to decision-tree mehods that construct intervals using the data provided).

The classic setting for constructing descriptions in logic from data is Inductive Logic Programming (ILP: [17]). ILP methods have often been applied to very small datasets. Most recently, it was shown that the ILP system DeepLog [19] was capable of generalising from even a single data instance, using a search method that preferred increasingly specific descriptions of the data. To the best of our understanding, DeepLog does not (as yet) handle continuous hypothesis-spaces and does not allow for noise in the data (both of which are needed here). We therefore propose a DeepLog-inspired approach that is tailored specifically for the problem we address in this paper.

Conceptually, it will be helpful for us to view ILP for molecule-generation as an experiment-guided search through a space of logical representation of feasible molecules (Fig. 2).

**Figure 2.**
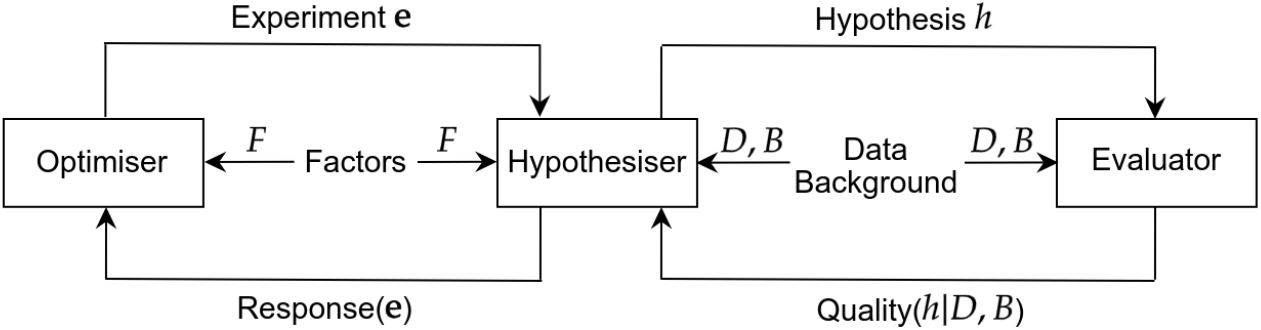
A hypothesize-and-test approach for multi-constraint molecule generation guided by an optimiser. *F* is some set of controllable numeric-valued factors and **e** denotes an *experiment* : an assignment of a range of values for each factor; *D* is a dataset of some labelled positive (“good”) and negative (“bad”) molecules; *H* is a symbolic hypothesis for feasible molecules; and *B* refers to background knowledge that includes any information about the target, and procedures needed to evaluate the quality of the generative hypothesis being proposed.

We clarify by way of example what we mean by a hypothesis for feasible molecules.

#### Example 1

(A Hypothesis About Feasible Molecules). *In this stage, we will be examining clausal descriptions of feasible molecules of this kind:*

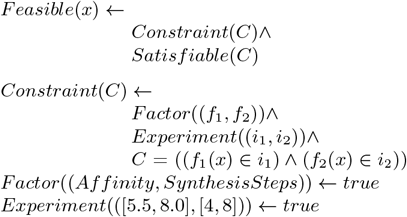

A complete formalisation of terms and concepts needed for the implementation can be found in Appendix A. Here we only include the two main definitions needed, and focus on some interesting consequences.

#### 2.1.1 The Hypothesiser and the Evaluator

Given a factor-specification and an experiment as defined in Appendix A, we can define the corresponding set of feasible instances declaratively as a clause. We will call this a *hypothesis*.

##### Definition 1

(Hypothesis). *Let 𝒳 be a set of instances*, (*F*, **Θ**) *be the factor specification and* **e** *an experiment given* (*F*, **Θ**). *Let X*_*F*,**e**_ *be the set of feasible instances given F*, **e**. *Let Satisfiable*(Φ_*F*,**e**_(*x*)) *is true if x* ∈ *𝒳 and x* ∈ *X*_*F*,**e**_. *Then a hypothesis is the clause:*

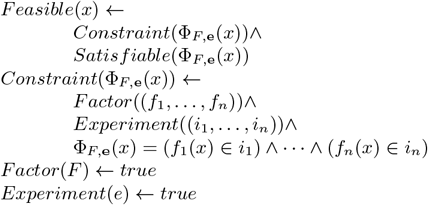

##### Example 2

(A Hypothesis About Molecules). *Suppose the controllable factors are Affinity and SynthesisSteps. A possible logical hypothesis is:*

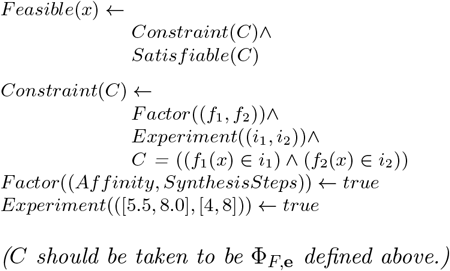

It is evident that the hypothesiser is simply a clausal representation of the set of feasible instances given a factor-specification and an experiment. The clausal form is not necessary, but is convenient since it allows us to directly use results from ILP on Bayesian-scoring of clausal hypotheses.

##### The Evaluator

The evaluator assesses hypotheses using the Bayesian *Q*-heuristic proposed by McCreath [20]. We motivate this with an example showing the inadequacy of simply using derivability of data instances.

###### Example 3

(Comparing Hypotheses). *Suppose we are given a dataset containing 1 example of a good molecule m*_1_ *(a “positive” example). Suppose we have 2 hypotheses:*

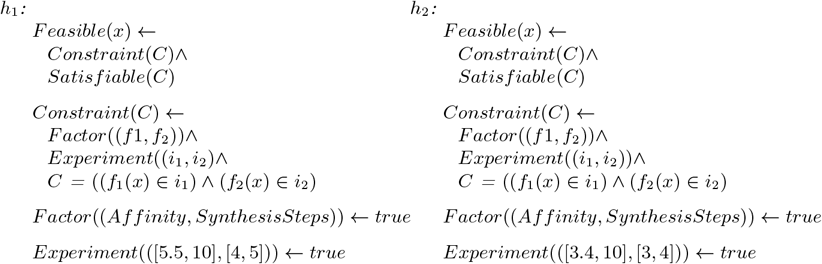

*Suppose we can derive Affinity*(*m*_1_) = 6.0 *and SynthesisSteps*(*m*_1_) = 4 *using the background knowledge B. Let us assume B* |= *Satisfiable*(*C*) *iff C has at least 1 model. Clearly B* ∧ *h*_1_ |= *Feasible*(*m*_1_) *and B* ∧ *h*_2_ |= *Feasible*(*m*_1_).

We are therefore unable to distinguish between *h*_1_ and *h*_2_ in the example, just given derivability of *Feasible*(*m*_1_). McCreath’s *Q*-heuristic is a Bayesian measure that captures tradeoffs between derivability, generality and complexity of a hypothesis (usually incorporated through the prior probability). It generalises the positive-only setting of [21], that is also used in DeepLog (McCreath allows for learning from positive or negative examples, and allows for noise in the labels). In the following *h*|*B* is to be read as “*h* given *B*”.

###### Definition 2

(McCreath’s *Q*-Heuristic). *Let 𝒳denote the set of all instances. Let h be a hypothesis as defined in Defn. 1. Let E*^+^ *denote a set of positive (feasible) examples and E*^−^ *denote a set of negative (infeasible) examples s*.*t*. (|*E*^+^| + |*E*^−^|) *>* 0. *Let D* = *E*^+^ ∪ *E*^−^. *Let ext*(*h*|*B*) = {*x* : *x* ∈ 𝒳, *B* ∧ *h* |= *Feasible*(*x*)}; *and for any* 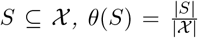. *Let ϵ be the probability that an instance is randomly assigned to E*^+^ *(resply. E*^−^*). Let B denote background knowledge, TP* (*h*\*B, D*) = {*e* : *e* ∈ *E*^+^, *e* ∈ *ext(h*|*B)*} ; *TN* (*h*|*B, D*) = {*e* : ¬*e* ∈ *E*^−^, *B* ∧ *h* ∧ *e ext(h*|*B)*} ; *and FPN* (*h*|*B, D*) = *D \* (*TP* (*h*|*B, D*) ∪ *TN* (*h*|*B, D*)). *Then, dropping the inclusion of B, D for convenience, the fixed-example model in McCreath [20] defines the quality of a hypothesis as:*

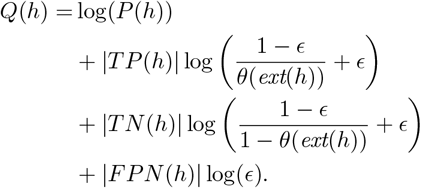

*For the special case of ϵ* = 0, *the quality of a hypothesis in the fixed-example setting simplifies to:*

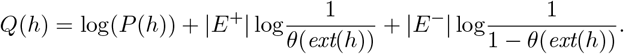

In [20] it is shown that maximising *Q*(*h*|*B, D*) maximises the Bayesian posterior P(*h*|*B, D*), along other theoretical results including a proof of (probabilistic) convergence to a target concept. Assuming the entailment relation |= can be checked, the practical difficulties in using the *Q*-heuristic are in obtaining the values for *θ*(*ext*(*h*)) and *P* (*h*). We note the following:

###### Remark 1.

*We will need the following to be able to use the Q-heuristic here:*

a. *In order to obtain the sets TP, TN and FPN we will require B to contain all the definitions needed to evaluate the constraint in the hypothesis (that is, B will need to contain definitions for the f*_*i*_(·)).
b. *By definition, ext*(*h*) *is the set of feasible instances as defined in Defn*.8 *in Appendix A. We can estimate θ*(*ext*(*h*)) *on a random sample X* ⊂ 𝒳 *as follows: Let S* = {*x* : *x* ∈ *X*, Φ(*x*) *is true*}. *Then the (maximum-likelihood) estimate of θ*(*ext*(*h*)) *is* 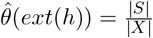.^2^

###### Example 4

(Comparing Hypothesis (again)). *For the hypotheses h*_*i*_ *in Example 3*, |*TP* (*h*_*i*_)| = 1, |*TN* (*h*_*i*_)| = |*FPN* (*h*_*i*_)| = 0. *Let ϵ* = 0 *and P* (*h*_1_) = *P* (*h*_2_). *Then:*

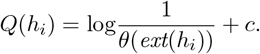

*If* (*ext*(*h*1) < *ext*(*h*2)), *then Q*(*h*1) > *Q*(*h*2).

More generally, we note the following:

###### Remark 2.

*These results follow straightforwardly from the definition in Defn. 2*:

**Positive-only data**. *Let E*^+^ ≠ ∅ *and E*^−^ = ∅. *Let ϵ* = 0. *If P* (*h*_1_) = *P* (*h*_2_), *then* (*Q*(*h*_1_) *> Q*(*h*_2_)) *iff* (*ext*(*h*_1_) *< ext*(*h*_2_)).

**Negative-only data**. *Let E*^−^ ≠ ∅ *and E*^+^ = ∅. *Let ϵ* = 0. *If P* (*h*_1_) = *P* (*h*_2_), *then* (*Q*(*h*_1_) *> Q*(*h*_2_)) *iff* (*ext*(*h*_1_) *> ext*(*h*_2_)).

*That is, in the noise-free case, with equal prior probabilities, and positive data only, more specific hypotheses will be preferred; and with negative data only, more general hypotheses will be preferred*.

### 2.2 The Optimiser

The goal of the optimiser is to identify the best hypothesis (measured by the *Q*-heuristic just described). `SearchHypothesis` (Procedure 1) shows the main steps of a simple greedy search procedure that, given a factor-specification: (a) Randomly samples experiments (hyper-rectangles) within a pre-defined boundary; (b) Evaluates the corresponding hypotheses; (c) Selects the experiment that results in the best hypothesis; and (d) Repeats from (a) with the selected experiment defining the boundary.

#### Example 5

(Nested Rectangles). *Suppose* `SearchHypothesis` *is attempting to find a hypothesis given 2 factors F* = (*Affinity, SynthesisSteps*), *with* Θ = *e*_0_ = ([5, 10], [4, 8]). *Then* `SearchHypothesis` *starts with the constraint* (*Affinity* ∈ [5, 10]) ∧ (*SynthesisSteps* ∈ [4, 8]), *which corresponds to a rectangle R*_0_ *in the Cartesian-space with Affinity and SynthesisSteps. Let the Q-value of the corresponding hypothesis be Q*_0_. `SearchHypothesis` *then randomly samples rectangles contained within R*_0_. *Suppose the rectangle R*_1_, *defined by e*_1_ = ([6, 10], [4, 6]), *and the corresponding hypothesis has the highest Q-value (Q*_1_*) of all the rectangles sampled*. `SearchHypothesis` *then iterates by sampling within R*_1_. *The search procedure therefore identifies a sequence of nested rectangles*.

We have not described how the sampling is done in Step 7 of `SearchHypothesis`. Here there are several options: the easiest to sample each dimension of the bounding hyper-rectangle independently, using a uniform distribution. Better sampling procedures exist (for example, Latin Hyper-rectangle Sampling [22], DIRECT [23], Bayesian sub-region sampling [24] and so on). Alternatives to the the simple greedy strategy of picking the best-scoring hyper-rectangle in Step 9 also also clearly possible, by drawing an experiment from the distribution of scores in Step 8.^3^

#### Procedure 1

`SearchHypothesis`

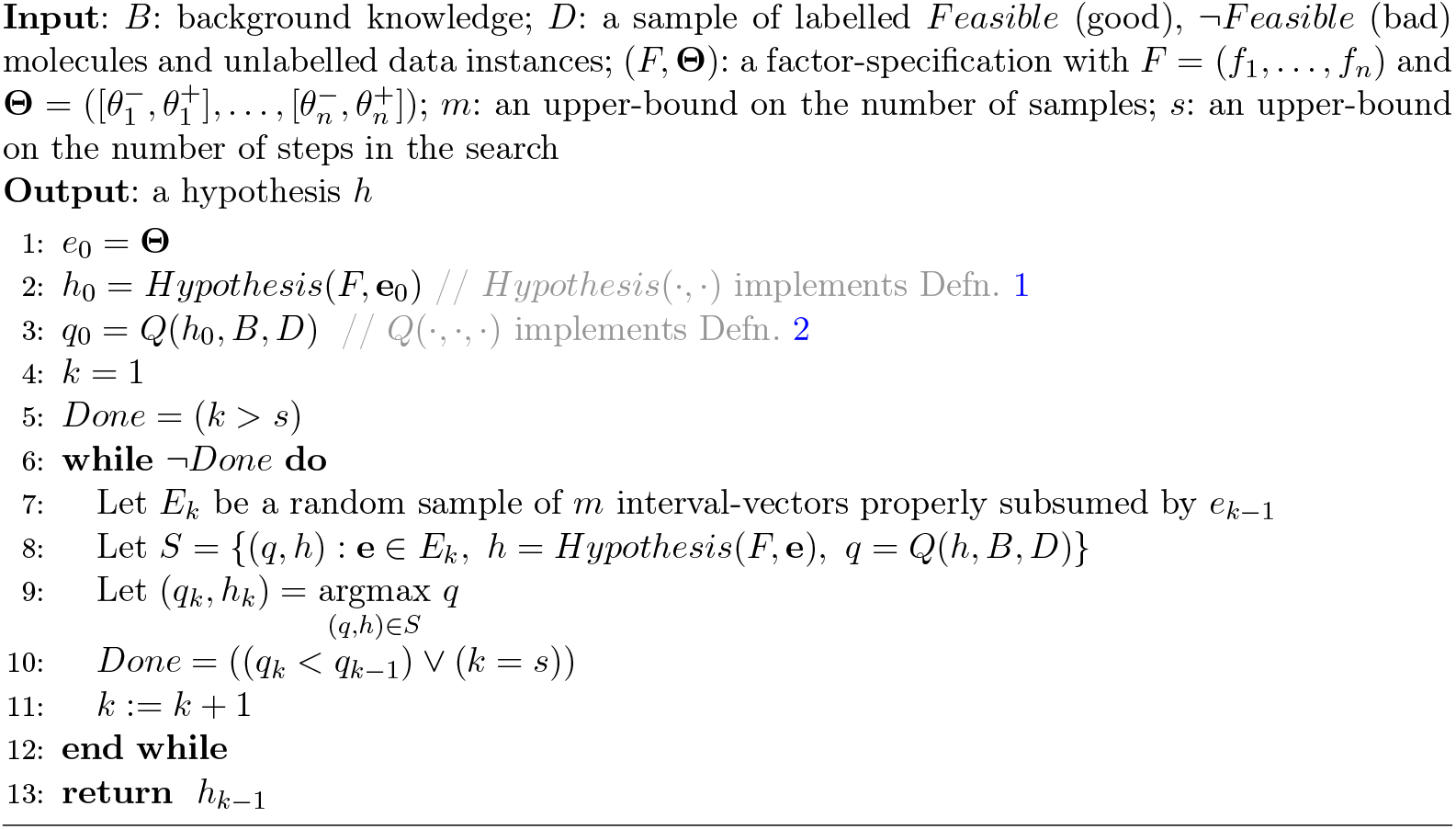

#### Example 6

(Sampling(Hyper-)Rectangles). *Suppose* `SearchHypothesis` *is attempting to sample a rectangle bounded by* [*x*_1_, *x*_2_] *and* [*y*_1_, *y*_2_]. *Examples of some strategies for sampling sub-rectangles are:*

#### Uniform orthogonal sampling

*(a) Select a pair of points a* = *U* (*x*_1_, *x*_2_) *and b* = *U* (*x*_1_, *x*_2_) *s*.*t. x*_1_ *< a < b < x*_2_; *(b) Select a pair of points c* = *U* (*y*_1_, *y*_2_) *and d* = *U* (*y*_1_, *y*_2_) *s*.*t. y*_1_ *< c < d < y*_2_; *and (c) The new rectangle is bounded by* [*a, b*] *and* [*c, d*].

#### Sampling with fixed upper- or lower-bounds

*(a) Select a point a* = *U* (*x*_1_.*x*_2_) *s*.*t. x*_1_ *< a < x*_2_; *(b) Select a point d* = *U* (*y*_1_, *y*_2_) *s*.*t. y*_1_ *< d < y*_2_; *and (c) The new rectangle is bounded by* [*a, x*_2_] *and* [*y*_1_, *d*].

We treat the choice of sampling method as an application-specific detail. The following property will however hold for any procedure that progressively selects (hyper-)rectangles subsumed by a bounding hyper-rectangle

##### Proposition 1.

*Let* (*F*, ·) *be a factor-specification, B denote background knowledge. Let* **e**_*k*_ (1 ≤ *k* ≤ *s) be an experiment selected by the iterative procedure* `SearchHypothesis` *on the k*^*th*^ *iteration s*.*t*. **e**_*k*_ *is subsumed by* **e**_*k*−1_. *Let h*_*k*_ = *Hypothesis*(*F*, **e**_*k*_), *and h*_*k*−1_ = *Hypothesis*(*F*, **e**_*k*−1_). *Then h*_*k*−1_ |= *h*_*k*_.

*Proof*. First we observe that *h*_*k*_ and *h*_*k*−1_ differ only in the constraint which appears in the body of (the unfolded form of) each clause and *Constraint*(Φ_*F*,**e**_(*a*)) is true for any *a*, **e**. Suppose there exists an element *a* of 𝒳 such that *h*_*k*−1_(*a*) is true but *h*_*k*_(*a*) is false. Hence *Feasible*(*a*) ← *Constraint* 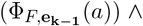 *Satisfiable* 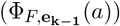 is true among other things and *Feasible*(*a*) ← *Constraint* 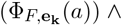 *Satisfiable* 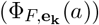 is false. It is only possible when both *Constraint* 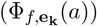 and *Satisfiable* (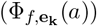 are true but *Feasible*(*a*) is false. *Satisfiable* 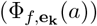 is true implies 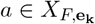. Since **e**_**k**−**1**_ subsumes **e**_**k**_, 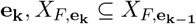 (refer Remark 1). So a is in 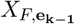. Hence *Satisfiable* 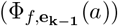 is true. Hence *Feasible*(*a*) is true which is a contradiction. So for all *a* ∈ 𝒳, whenever *h*_*k*−1_(*a*) is true then *h*_*k*_(*a*) is also true. ■

##### Remark 3.

*In the terminology of Inductive Logic Programming (ILP), h*_*k*−1_ |= *h*_*k*_ *means h*_*k*_ *is more specific than h*_*k*−1_. `SearchHypothesis` *therefore examines progressively more specific hypotheses*.

In that next remark, we conclude that ext(*h*_*k*_) ⊆ ext(*h*_*k*−1_) using the above proposition and proposition 3.

##### Remark 4.

*Let* (*F*, ·) *be a factor-specification, B denote background knowledge. Let* **e**_*k*_ (1 ≤ *k* ≤ *n) be an experiment selected by an iterative procedure on the k*^*th*^ *iteration s*.*t*. **e**_*k*_ *is subsumed by* **e**_*k*−1_. *Let h*_*k*_ = *Hypothesis*(*F*, **e**_*k*_), *and h*_*k*−1_ = *Hypothesis*(*F*, **e**_*k*−1_). *Then ext*(*h*_*k*_) ⊆ *ext*(*h*_*k*−1_).

Finally, we comment on the relation to the constraints constructed by the LMLF technique in [1].

##### Remark 5

(Relation to LMLF Hypothesis Space). *Suppose we are only interested in numeric factors. We note informally that: Every conjunct in [1] can be mapped to a corresponding hypothesis here. This follows straightforwardly from the fact that every hypothesis in [1] can be rewritten in the form defined in Defn. 1, provided interval bounds are known for each numeric factor. Suppose C is a conjunct in [1]. Then, assuming the bounds* [*a*_*i*_, *b*_*i*_] *for each factor f*_*i*_ *the constraint C*^*′*^ *the body of the hypothesis according to Defn. 8 is obtained as follows. if* (*f*_*i*_(*x*) ≤ *θ*_*i*_) *is a term in C then* (*f*_*i*_(*x*) ∈ [*a*_*i*_, *θ*_*i*_]) *is a term in C*^*′*^; *if* (*f*_*i*_(*x*) ≥ *θ*_*i*_) *is a term in C then* (*f*_*i*_(*x*) ∈ [*θ*_*i*_, *b*_*i*_]) *is a term in C*^*′*^; *otherwise* (*f*_*i*_(*x*) ∈ [*a*_*i*_, *b*_*i*_]) *is in C*^*′*^.

*Thus the hypothesis space considered by LMLF is a subset of the hypothesis space considered here. Differences also arise in the search procedures used. LMLF requires the background knowledge to specify the θ*_*i*_ *above. Here, these are automatically determined by the sampling procedure and greedy search using the Q-heuristic*.

### 2.3 Stage 2: Logical Specification to Program

Given a factor-specification (*F*, ·), the hypothesis obtained from Procedure 1 is a clause:

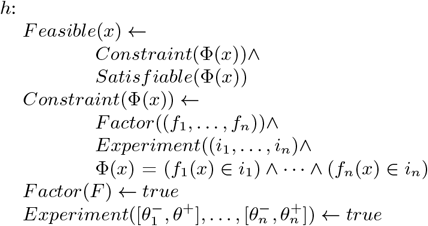

Along with suitable definitions in the background knowledge, this clause can be used for checking feasibility, but it does not give us a program for generating new molecules. We propose to achieve this by adapting the generative part of the procedure described in [1]. This part is an iterative procedure that repeatedly: (a) generates instances using a large language model (LLM); (b) uses the symbolic hypothesis identified along with definitions in the background knowledge to test if the instances are feasible; (c) Updates the context of the LLM with the result of the constraint-satisfaction test; and (d) repeats from (a). A program is shown in Procedure 2, Ignoring various bounds, it is evident that the set of molecules obtained is the result of the generalised composition `LMLFStar` (*λ, B*,`SearchHypothesis` (*B, D*, (*F*, **Θ**))). The reader would have anticipated a problem with decomposing of the discriminative step (done by `SearchHypothesis`) and generative step (done by `LMLFStar`) in this manner. We will address this below.

#### Proposition 2

(Correctness). *Let* 𝒳 *denote the set of all molecules; and let* (*B*∧*h*) *be consistent. Let ext*(*h*|*B*) = {*x* : *x* ∈ 𝒳, (*B* ∧ *h*) |= *Feasible*(*x*)} *denote the subset of* 𝒳 *identified as feasible by* (*B* ∧ *h*). *Then the set M*_*n*−1_ *returned by* `LMLFStar` *is a subset of ext*(*h*|*B*).

*Proof*. `LMLFStar` executes the loop (Steps 4–11) at most *n* times. We claim the following is loop invariant:

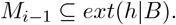

#### Procedure 2

`LMLFStar`

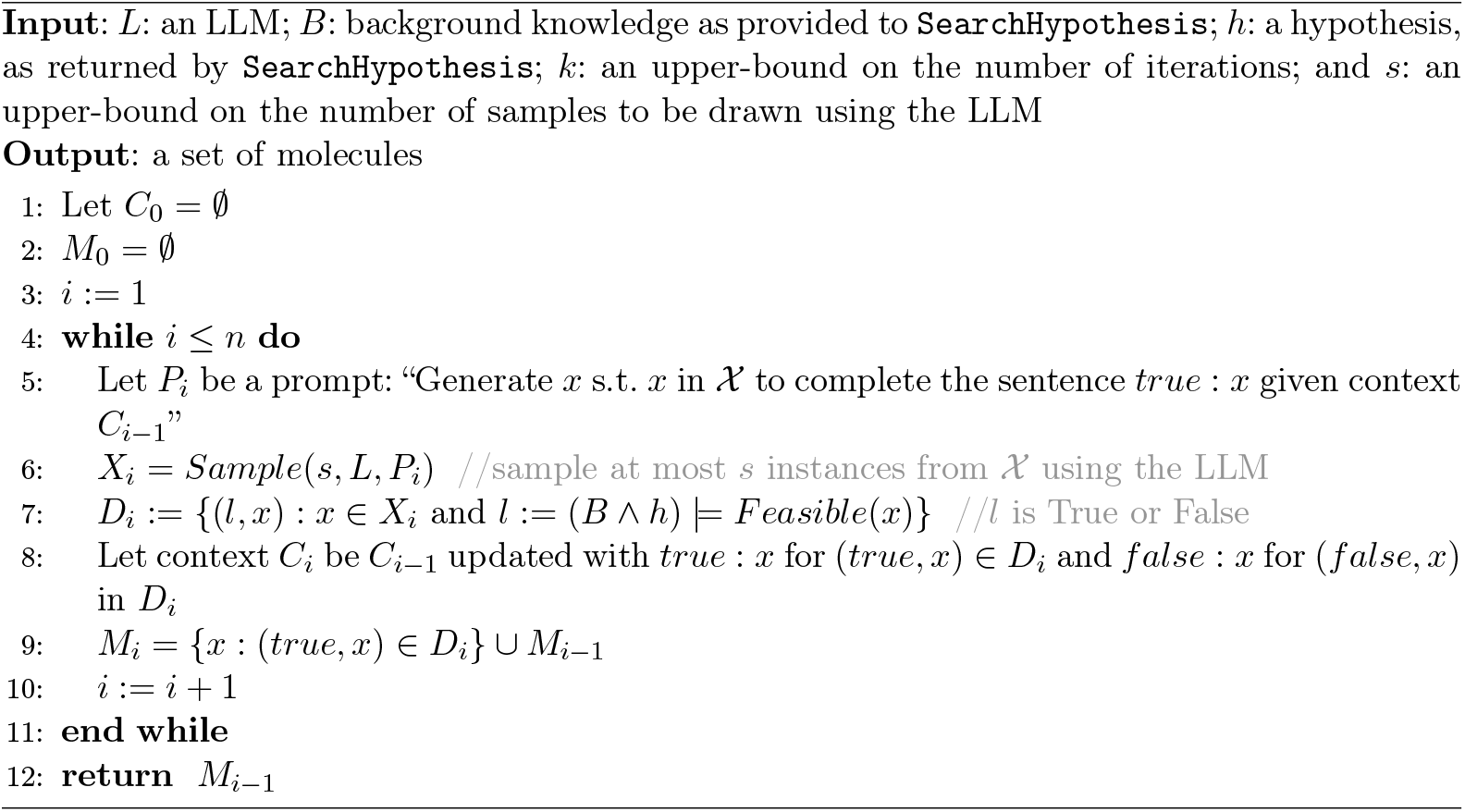

At the start of the iteration (*i* = 1), *M*_*i*−1_ = ∅, and the invariant is trivially true. Assume the invariant holds at the start of the *k*^th^ iteration. That is, *M*_*k*−1_ ⊆ *ext*(*h*|*B*). In Step 9 *M*_*k*_ = *X* ∪ *M*_*k*−1_, where *X* = {*x* : *x* ∈ 𝒳, (*true, x*) ∈ *D*_*k*_}. Since (*true, x*) ∈ *D*_*k*_ iff (*B* ∧ *h*) |= *Feasible*(*x*), *X* ⊆ *ext*(*h*|*B*). That is, *M*_*k*_ ⊆ *ext*(*h*|*B*). The loop variable *i* is incremented to *k* +1, and at the start of the next iteration, clearly *M*_*i*−1_ = *M*_*k*_ ⊆ *ext*(*h*|*B*). The procedure clearly terminates since *i* is bounded by *n*, and the procedure returns the set *M*_*n*−1_ which is a subset of *ext*(*h*|*B*). That is every molecule of the set returned by `LMLFStar` is a feasible molecule as defined by (*B* ∧ *h*). ■

We now turn to the sequential decomposition of discriminative and generative steps. The greedy approach in `SearchHypothesis` can result in a hypothesis that is overly-specific, resulting in `LMLFStar` returning an empty set of molecules. The “fix” is to interleave the two steps. An implementation is shown in Procedure GenMol. The reader can verify that `GenMol` is largely the same as `SearchHypothesis`: the difference is that calls to the generator are made as the search proceeds.^4^

## 3 Empirical Evaluation

We consider two kinds of experiments:

### Validate

We evaluate the performance of `GenMol` in the controlled setting examined in [1], with a known target-site, a large number of known inhibitors and non-inhibitors, and a single factor to be optimised (estimated binding affinity to the target-site).

### Explore

We evaluate the performance of `GenMol` in an open-ended setting where the true target-site is not known precisely, and multiple factors have to be optimised (binding affinity, molecular weight and synthesis accessibility), and there are very few known inhibitors.

#### Procedure 3

`GenMol`

In the first category, our assessment will be largely statistical. In the second category, we will also obtain the evaluation of a synthetic chemist.

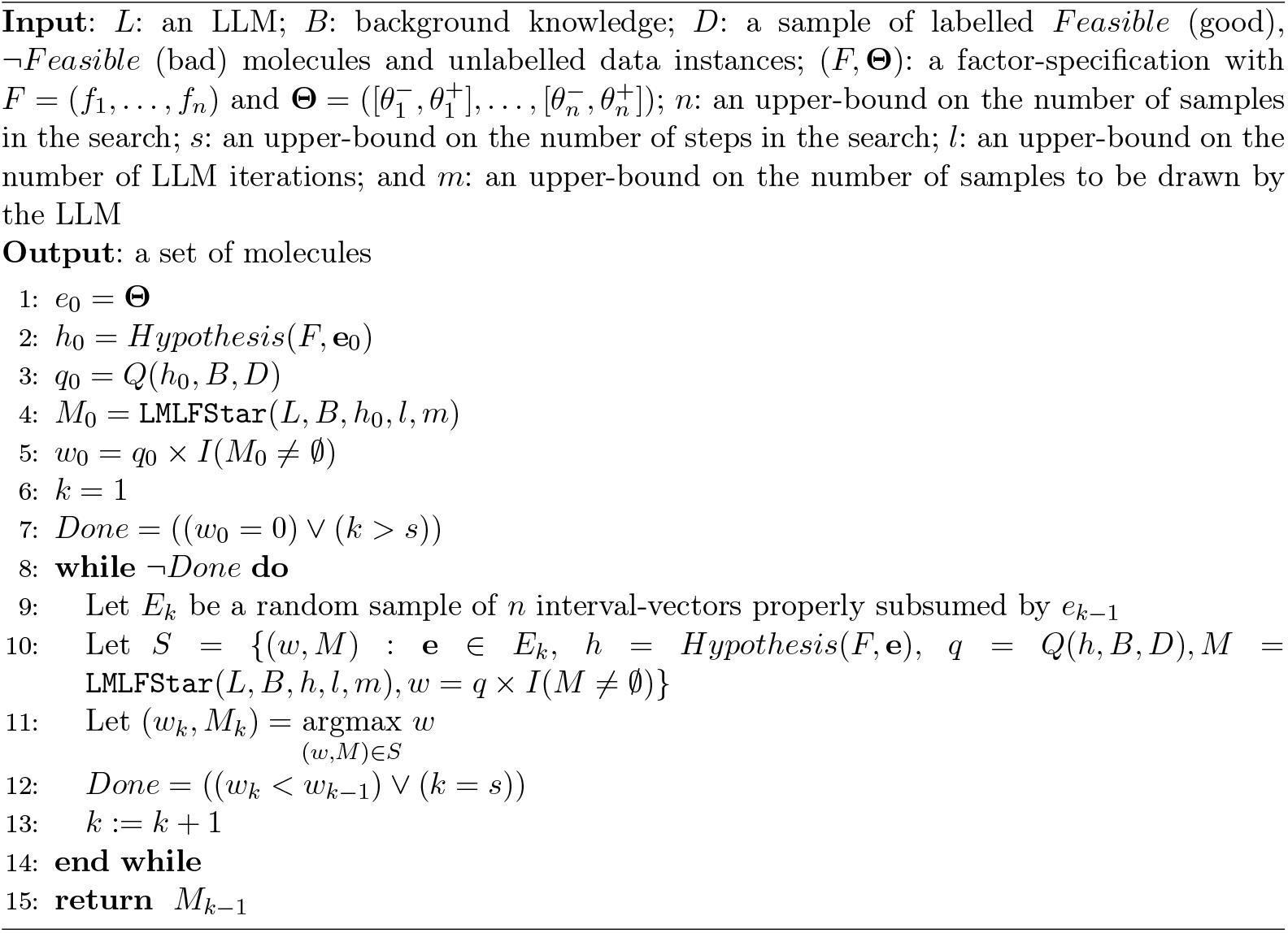

### 3.1 Materials

#### 3.1.1 Problems

##### Kinase Inhibitors

We conduct our controlled evaluations on 2 well-studied kinase inhibitors: (a) JAK2, with 4100 molecules provided with labels (3700 active); and (b) DRD2 (4070 molecules with labels, of which 3670 are active). These datasets are from the ChEMBL database [25], which are selected based on their *IC*_50_ values and docking scores. JAK2 inhibitors are drugs inhibit the activity of the Janus kinase 2 (JAK2) enzyme, in turn affecting signalling pathways, especially to the cell nucleus. These pathways are critical for various immune response reactions and are used to develop drugs for autoimmune disorders like ulcerative colitis and rheumatoid arthritis. DRD2 (dopamine D2) inhibitors are drugs that block dopamine”s ability to activate the DRD2 receptors. This reduces dopamine signalling, and is used to treat psychological disorders like schizophrenia.

##### DBH Inhibitors

Human dopamine *β*-hydroxylase (DBH), is an enzyme that converts dopamine (DA) to norepinephrine (NA), plays a pivotal role in regulating the concentration of NA deficiency or overproduction of which causes several diseases related to the brain and the heart. This enzyme is thus of high therapeutic significance. The availability of the three-dimensional structures of DBH is expected to facilitate the identification of DBH active-site inhibitors. In the meantime, the crystal structure of a dimer of DBH has been determined, providing insights into its function and aiding in the design of inhibitors [26]. Specifically, we will use the *in silico* model of the dimer generate small molecules with similar or better IC50 and KD values (in simulation) than at least one of the latest generation DBH inhibitor. We will use as data the 5 known DBH inhibitors: Tropolone, Disulfiram, Nepicastat, Zamicastat, Etamicastat. The last three are shown in Fig. 3. Tropolone is a naturally occurring molecule, with known toxic effects. Disulfiram is a 1st generation molecule, and also with toxic side-effects. The last two molecules, Zamicastat and Etamicastat are the latest generation of DBH inhibitors and are currently in double-blind human trials for hypertension. We focus on obtaining molecules with docking scores at least as good as Nepicatat, a 4th generation drug.

**Figure 3.**
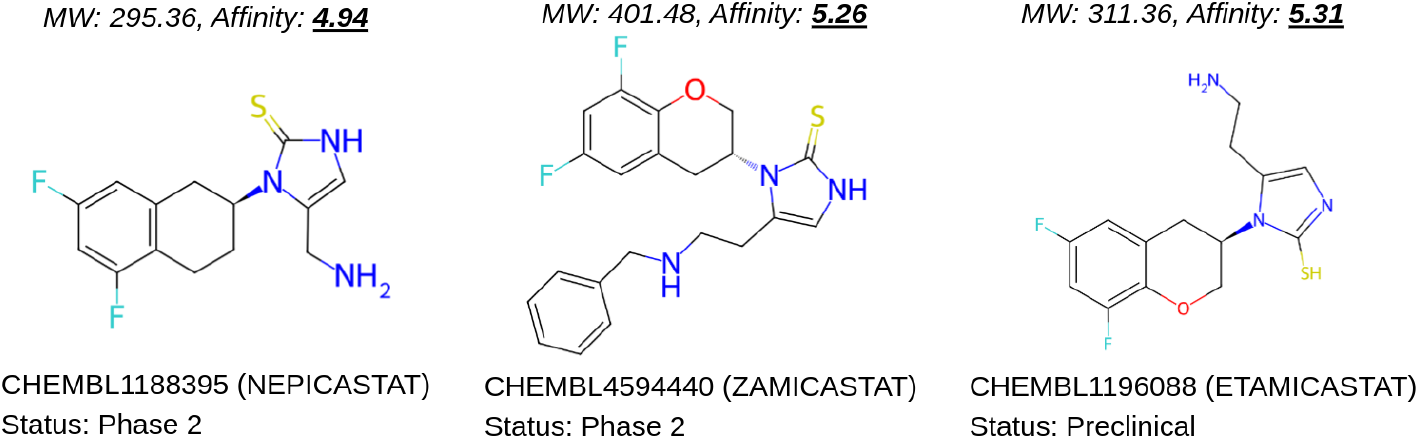
Three known DBH inhibitors at different stages of their FDA approval status. Each compound is identified by their CHEMBL ID and name. “MW” refers to molecular weight; “Affinity” refers to the binding affinity predicted by `GNINA` software while docking the molecules to the DBH protein, *4zel*. The approval status of these compounds were noted as on 25 January 2025 from the ChEMBL online portal.

#### 3.1.2 Background Knowledge

We distinguish the following components of background knowledge:

A. Specialists” knowledge. We require biological knowledge of the target site, or a proxy for the target size. We also need chemical knowledge of the relevant factors, the range of their values, and information (if any) of whether the factors need to be maximised or minimised.
B. Factor-functions. These are definitions for computing the factors (like *Affinity, MolWt etc*.) for molecules. Usually this will also include procedures provided by some molecular modelling software (like RDKit).

#### 3.1.3 Algorithms and Machines

All the experiments are conducted using a Linux (Ubuntu) based workstation with 64GB of main memory and 16-core Intel Xeon 3.10GHz processors. All the implementations are in Python3. We use `OpenAI` library (version 1.52.1) for sampling molecules from `GPT-4o` [27]. We use `RDKit` (version 2024.09.3) [28] for computing molecular properties and `GNINA` (version 1.3) [29] for computing docking scores (binding affinities) of molecules. Additional details of the experimental setup can be found in Appendix B. We have used PubChem Sketcher V2.4 for drawing the 2-D structures of the molecules shown in this paper.

### 3.2 Method

Our method for both controlled and open-ended experiments is straightforward and follows these steps:

1. Identify factor-set *F* and other bounds.
2. Obtain data instances consisting of positive, negative and unlabelled examples. Any positive example is taken to be feasible and any negative example is taken to be infeasible.
3. Using the background knowledge *B* described in Sec. 3.1, *D, F* and other bounds:
  a. Obtain a set of molecules using `GenMol`;
  b. Assess the quality of the molecules generated.

The following additional details are relevant:

- In the controlled (Validate) experiments, we are only attempting to optimise one factor, namely: docking score, which is indicative of binding affinity. This is in line with what was done in [1]. For the open-ended (Explore) experiments, we will extend this to include: number of synthesis steps, and estimated yield per step.
- The factor-set specification also requires identifying minimum and maximum for the initial search space. For all experiments, we use: {*affinity* : [3, 10], *molwt* : [200, 700], *SAS* : [0, 7.0]}, where *affinity* is the predicted affinity from GNINA software, *molwt* is the molecular weight, and *SAS* denotes synthesis accessibility score.
- For the controlled experiments, we sample 30 inhibitors and 30 non-inhbitors to be part of the dataset *D*. For the open-ended experiments we use the 5 molecules tabulated in Sec. 3.1, and will take Tropolone as a non-inhibitor. Estimating the *Q*-heuristic requires a sample of unlabelled molecules. For this we use a randomly drawn set of 1000 molecules from the ChEMBL database.
- The description of `SearchHypothesis` does not specify a sampling method for obtaining subsumed hyper-rectangles. We use an approach based on Latin Squares Hyper-Rectangle Sampling (LHRS: [22]). Some additional prior information may be available that may be used to modify the basic LHRS approach: (a) If we know beforehand that a factor is to maximised (for example, binding affinity), then we do not sample points from the upper-end of the range for the factor. Thus, subsuming rectangles are obtaining by only random placements of the lower-end; (b) Similarly, if we know beforehand that we want to minimise a factor, then only the upper-threshold is sampled. If nothing is known then a standard LHRS approach is adopted.
- For all experiments, we use a value of *s* = 10; and *n* = 10 for `GenMol` and we sample 100 molecules when it returns the optimal hypothesis. The set of feasible molecules during search and in the final generation are considered for evaluation.
- We assess the results of controlled experiments by examining the range and median docking scores of the molecules generated, and compare those to those generated by the LMLF procedure in [1]. For the open-ended experiments, we obtain the statistics on docking scores, and compare them to those of the latest generation of molecules used for DBH inhibition (see Sec. 3.1). In addition, we also provide an assessment of the molecules by an expert synthetic chemist.
- For each target problem, we assess the novelty of the generated molecules by using the average Tanimoto (or Jaccard) coefficient to the database of known inhibitors.

### 3.3 Results

#### Validate

Figure 4 a comparison against the results tabulated for LMLF++ in [1]. LMLF++ was the best performing variant in that paper, and was subtantially better than previous benchmarks set by the use of a VAE-GNN combination and reinforncement-learning based methods. It is evident from the tabulation that we are able to perform at least as well as LMLF++. It is also helpful if a leag-generator proposes novel molecules. For the JAK problems, this is not easy, since the number of known inhibitors is very large. Nevertheless, Fig. 5 sugests that the molecules generated by `GenMol` may still be quite novel (this is due to the prompt used for the LLM that attempts to generate molecules not in any known chemical database).

**Figure 4.**
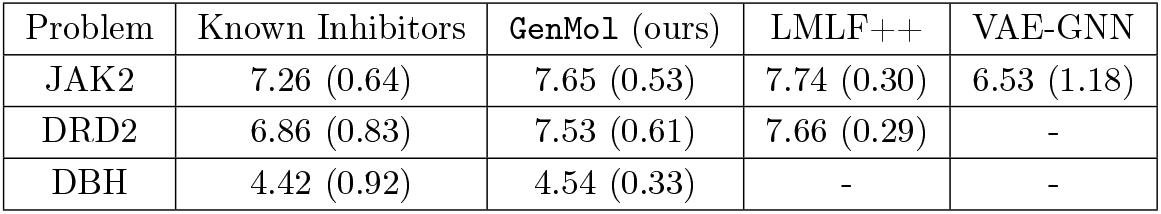
Statistics of binding affinities (the higher the better) for molecules obtained from `GenMol` on benchmark datasets. The entries represent the mean values, with standard deviations shown in parentheses. We compare against recent results using LMLF++ [1] and prior results using a VAE-GNN model [11]. “-” denotes “not available”.

**Figure 5.**
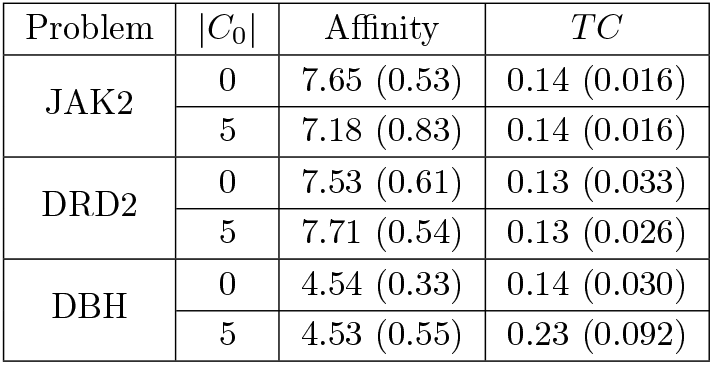
Statistics of binding affinities and novelty of LLM-generated molecules using `GPT-4o`. The entries represent the mean values, with standard deviations shown in parentheses. *C*_0_ denotes a set of known inhibitor of the target protein, provided as an early context to the LLM during prompting (see Appendix for more information). Predicted affinity refers to the predicted binding affinity from `GNINA` software. The predicted affinity is always non-negative, with higher values indicating stronger binding to the target protein. The Tanimoto coefficient (*TC*) ranges from 0 to 1, where 1 signifies highly similar structures, while 0 indicates complete structural dissimilarity.

#### Explore

The exploratory problem concerns generating potential leads for DBH inhibition, given data on the structure and inhibitory values of 5 molecules on a proxy target to DBH. The inhibitory efficacy of a small number of molecules is known (see Sec. 3.1). We consider two kinds of exploratory experiments. First, the LLM is provided with the information about the known molecules and their structure: in LLM parlance, we are doing “few-shot learning” (correctly, we are using the LLM to draw from a distribution conditioned on the known molecules). We will evocatively call this “In-the-Box” exploration. Secondly, we do not give the LLM any information about known molecules (“zero-shot learning”, or “Out-of-the-Box” exploration). The top-5 molecules obtained for each kind of experiment is shown in Fig. 6.

**Figure 6.**
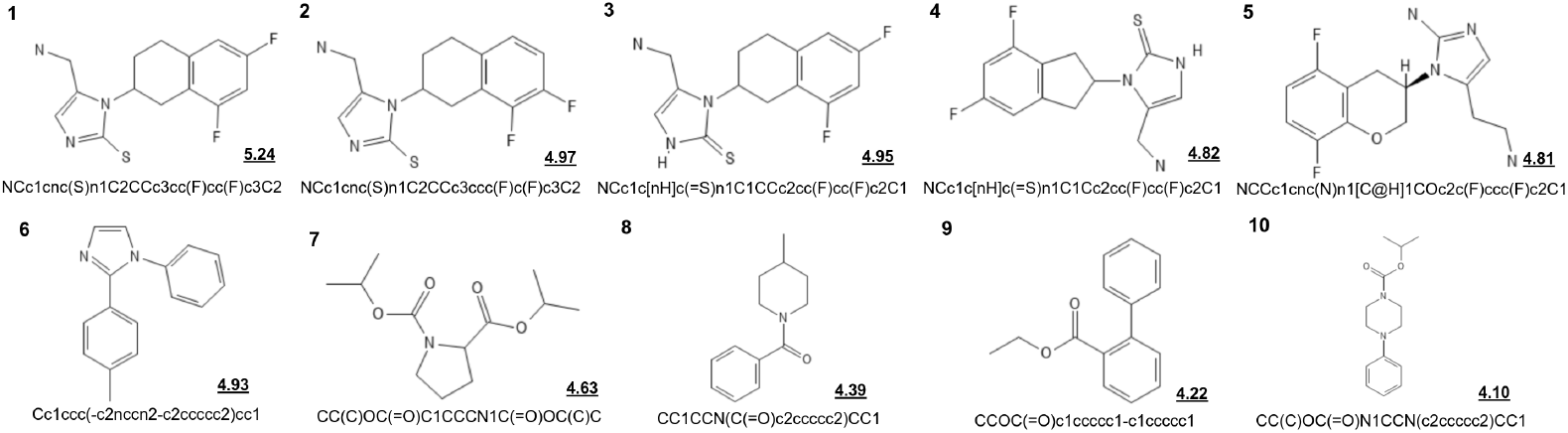
Potential inhibitors for DBH proposed by `GenMol`, along with their binding score to *4zel* protein. Molecules 1–5 are the top-5 molecules (ordered by estimated affinity) from “In-the-Box” exploration. Molecules 6–10 are from “Out-of-the-box” exploration. In the former the LLM used by `GenMol` has few-shot examples when it starts. In the former, no such information is provided, and LLM uses its underlying distribution over molecules are used to generate the molecules.

But are the molecules any good? Here are some assessments by specialists:

> **“In-the-Box” Exploration**. The views of the specialists about Molecules 1–5 are as follows:
>
> **Structural Biologist**. Molecules 1–4. These are [likely to be] good inhibitors of DBH since these molecules are structurally similar to dopamine and nepicastat. Another good feature of these molecules is that it carries Fluorenes as well. Molecule 5. Could be a better inhibitor of DBH since it is structurally similar to dopamine and nepicastat. It may work. It could be better than the four other molecules since it carries one oxygen in the aromatic ring. Another good feature of this molecule is that it also carries Fluorenes.
>
> **Synthetic Chemist**. All molecules can be synthesized, but is likely to be a long synthesis, and costs will be high.
>
> **“Out-the-Box” Exploration**. The views of the specialists about Molecules 6–10 are as follows:
>
> **Structural Biologist**. Molecule 6. This is a novel inhibitor of DBH where halogen is absent, OH/O is also absent. Not similar to dopamine or nepicastat. Thus, it can be an allosteric inhibitor of DBH. Molecules 7,8. This is also a novel inhibitor of DBH where halogen is absent. Since, O is present twice, the affinity might be tighter. Aromatic rings are very dissimilar to dopamine or nepicastat. Thus, mode of inhibition is difficult to predict. Molecule 9. May not be a good inhibitor due to highly flexible sidechains. Molecule 10. This can be a better inhibitor than the first one since its aromatic rings are very interesting to bind in the active site of DBH. O= is present and thus the affinity might be tighter. Mode of inhibition should be competitive.
>
> **Synthetic Chemist**. All molecules can be synthesised, and synthesis is likely to be through a short route. The molecules may be commercially available.

The broad takeaways are these: (a) Unsurprisingly, In-the-Box exploration appears to yield molecules that are similar (but not the same) as existing inhibitors. In contrast, it is very interesting that Out-of-the-Box exploration appears to yield very different molecules to known inhibitors; (b) There are good biological reasons to expect Molecules 1–5 to bind to the target, and Molecules 6,7,8 and 10 to bind to the target. Of these, the biologist believes Molecules 6 and 10 to be especially interesting; and (c) On the synthesis side, all molecules appear to be synthesisable, but 6–10 appear to be more amenable to a short (and therefore, possibly cheaper route).

It is noteworthy that biologists and chemists are able to comment meaningfully on instances generated by the machine. Additionally, not included here for reasons of space is a further round of ‘molecule-exchange’ between the chemist and the biologist, inspired by Molecules 1–5. The chemist proposed edited versions with shorter synthesis steps, and the biologist commented further on the biological suitability of those molecules. It is encouraging to see `GenMol`’s output is sufficiently intelligible and interesting to specialists to allow such interactions.

## 4 Concluding Remarks

This paper is concerned with the use machine learning (ML) techniques to accelerate the identification of ‘leads’ in early stage drug-design. Leads are small molecules capable of binding to a target protein, and satisfying some physico-chemical constraints. The specific problem we have examined is this:

**Given:** (a) information about of the target from a structural biologist; and (b) requirements on physico-chemical factors from a synthetic chemist.

**Find:** A program for generating a (potentially novel) set of molecules consistent with the specialists’ requirements.

Inspired by formal program design, we have approached this in two stages: first, constructing a logical specification, and the secondly, using the specification to obtain a program. Unlike with classic formal methods, ML techniques play a role in both stages. Inductive Logic Programming (ILP) is used to identify the logical specification from (possibly very small numbers of) data instances relevant to the target, and LLM technology is used to generate molecules consistent with the specification.

Conceptually, the principal advantage of decomposition into specification and implementation allows us to deal with with each in a modular way. Practically, this means tje effects of changes in specialists’ requirements or ML technology (for example, more appropriate ILP or LLM engines) can be localised. If we accept that this decomposition is useful, and that we do not want to construct the individual components entirely manually then the use of ILP and LLMs becomes almost obvious. The former has a long history of being able to identify logical descriptions even from very small datasets,7 given domain-knowledge. The latter is rapidly becoming the technique of choice for generating strings of all kinds, including molecules.

We have reported results from a validation study that shows that the approach works well on classic benchmark targets. However, it is the exploratory study that we think is substantially more relevance to the area of early-stage drug-design. Specifically, it shows: (a) the use of very small numbers of data instances (5, in this case); (b) that the output from the generator program can be directly understood and criticised by specialists; and (c) some of the results from the generator–especially in the “Out of the Box” mode–may be biologically novel and can cost-effective to synthesise. This suggests that for at least two human specialists, the result is an example of ‘Strong Machine Learning’ in the sense identified by Michie [30]. While the output has been made intelligble to the specilst through the use of an LLM, verifying the LLM’s output has been made possible through the use of ILP. We believe this kind of neural-symbolic ML will play a play an increasingly important role in the design of human-machine collaborative systems.

## Code and Data Availability

The data and code are available at: https://github.com/tirtharajdash/LMLFStar.

## Acknowledgements

AS is a visiting Professorial Fellow at UNSW, TCS Affiliate Professor, and a member of the Anuradha and Prashant Palakurthi Centre for AI Research (APPCAIR) at BITS Pilani. This research is partly supported by: DBT project BT/PR40236/BTIS/137/51/2022 “Developing Predictive Models for ‘druglikeness’ of small molecules”; and CDRF project C1/23/184 “Silicon-to-Lead: AI-Driven Design, Synthesis and Development of New Drugs to Combat Cardiovascular Diseases”. The authors would like to acknowledge Aaron Rock Menezes for his implementation of the PyLMLF algorithm reported in [1]. The authors sincerely thank Professor Suman Kundu and Professor Sumit Biswas for their insightful discussions on DBH.

## A Definitions, Results and Observations

### Definition 3

(Interval-Vectors). *An n-dimensional interval-vector* **v** = ([*a*_1_, *b*_1_], …, [*a*_*n*_, *b*_*n*_]) *is an element of* (ℝ × ℝ)^*n*^ *where a*_*i*_ ≤ *b*_*i*_ *for i* ∈ {1, …, *n*}.

We will sometimes denote (ℝ × ℝ) as ℐ and (ℝ × ℝ)^*n*^ as ℐ^*n*^. The set ℐ^*n*^ is therefore the set of *n*-dimensional hyper-rectangles, and an interval-vector is a hyper-rectangle.

### Definition 4

(Interval-Vector Subsumption). *Given interval-vectors* **v**_1_, **v**_2_ ∈ ℐ^*n*^ *If* (**v**_2_[1] ⊆ **v**_1_[1]) ∧ · · · ∧ (**v**_2_[*n*] ⊆ **v**_1_[*n*]) *then we will say* **v**_2_ *is subsumed by* **v**_1_ *(resply*. **v**_1_ *subsumes* **v**_2_*) We denote this by* **v**_2_ ⊑ **v**_1_ *(resply*. **v**_1_ ⊒ **v**_2_*). If there exists at least one j* ∈ {1, …, *n*} *s*.*t*. **v**_2_[*j*] ⊂ **v**_1_[*j*], *we will say* **v**_2_ *is properly subsumed by* **v**_1_ *(resply*. **v**_1_ *properly subsumes* **v**_2_*). We denote this by* **v**_2_ ⊏ **v**_1_ *(resply*. **v**_1_ ⊐ **v**_2_).

Clearly, if **v**_1_ ⊏ **v**_1_ then **v**_1_ ⊑ **v**_2_. We are typically interested in a set of factors {*f*_1_, *f*_2_, …, *f*_*n*_}. It is convenient to assume a total ordering *<* over the factors, and represent it by the sequence (*f*_1_, …, *f*_*n*_). A full specification of factors requires more than just their definitions:

### Definition 5

(Factors). *Let* 𝒳 *be a set of instances. A factor is a function f* : 𝒳 ↦ ℝ.

### Definition 6

(Factor Specification). *Let F* = (*f*_1_, …, *f*_*n*_) *be a sequence of factors. A factor specification is the pair* (*F*, **Θ**) *and* 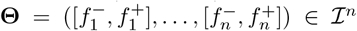. *For i* ∈ {1, …, *n*}, 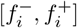 *is the range of values for the factor f*_*i*_.

A factor-specification allows us to define the notion of an *experiment* :

### Definition 7

(Experiment). *Let 𝒳 be a set of instances. Let* ((*f*_1_, …, *f*_*n*_), **Θ**) *be a factor-specification. An experiment* **e** *given the factor-specification, or simply an experiment, is an interval-vector in* ℐ^*n*^ *s*.*t*. **Θ** *subsumes* **e**.

An experiment imposes logical constraints on instances in 𝒳 and identify a set of *feasible instances* of 𝒳 for which the constraints are satisfied.

### Definition 8

(Feasible Instances). *Let* 𝒳 *denote a set of instances. Let* (*F*, **Θ**) *be a factor specification and* **e** *=* ([*a*_1_, *b*_1_], …, [*a*_*n*_, *b*_*n*_]), *be an experiment given F*, **Θ**. *Let* Φ_*F*,**e**(*x*)_ *be the boolean function* 𝒳 ↦ {*True, False*} *such that* Φ_*F*,**e**_(*x*) = ((*f*_1_(*x*) ∈ [*a*_1_, *b*_1_]) ∧ · · · ∧ (*f*_*n*_(*x*) ∈ [*a*_*n*_, *b*_*n*_])) *for x* ∈ 𝒳. *Then the set of feasible instances of* 𝒳 *given F, e, or simply the set of feasible instances, is the set X*_*F*,**e**_ = {*x* : *x* ∈ 𝒳, Φ_*F*,**e**_(*x*) = *true*}.

We note the following relationship between factor-specifications, experiments and feasible instances:

### Remark 6.

*Let* X *be a set of instances. Let* (*F*, **Θ**) *be a factor-specification and* **e**_**1**,**2**_ *be experiments given* (*F*, **Θ**). *If* **e**_**1**_ ⊒ **e**_**2**_ *then* 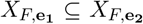. *This follows from the definition of the* ⊒ *relation (Defn. 4) and logical constraint defining feasible instances in Defn. 8*.

### Proposition 3.

*Let* (*F*, ·) *be a factor-specification, B denote background knowledge. If h*_1_ |= *h*_2_ *then ext*(*h*_2_) ⊆ *ext*(*h*_1_).

*Proof*. Suppose *x* is in *ext*(*h*_2_). So *B* ∧ *h*_2_ ⊧ *Feasible*(*x*). It is easy to see *B* ∧ *h*_1_ ⊧ *B* ∧ *h*_2_ using *h*_1_ ⊧ *h*_2_. Using the above two claims, we conclude *B* ∧ *h*_1_ ⊧ *Feasible*(*x*). Hence *x* ∈ *ext*(*h*_1_). So *ext*(*h*_2_) ⊆ *ext*(*h*_1_). ▪

### Remark 7

(Convexity of *Q*).

*Positive-only data. Let E*^+^ ≠ ∅ *and E*^*−*^ = ∅. *Let ϵ* = 0 *and h be a hypothesis s*.*t*. |*FPN* (*h*) | = 0, *and P* (*h*_1_) = *P* (*h*_2_) *for every h*_1_, *h*_2_. *Then Q*(*h*) *is a convex function*.

*Negative-only data. Let E*^*−*^ ≠ ∅ *and E*^+^ = ∅. *Let ϵ* = 0 *and h be a hypothesis s*.*t*. |*FPN* (*h*) | = 0, *and P* (*h*_1_) = *P* (*h*_2_) *for every h*_1_, *h*_2_. *Then Q*(*h*) *is a convex function*.

*Error-free Data. When ϵ* = 0 *and P* (*h*_1_) = *P* (*h*_2_) *for every h*_1_, *h*_2_, *Q*(*h*) *is a convex function. Q may not be convex in general*.

### Remark 8

(PAC Learnability). *Though it is not of practical relevance to the problems considered here, the class of axis-aligned rectangles exhibits interesting properties. For instance, the space of axis-aligned rectangles are shown to be efficiently learnable in the Probably Approximately Correct (PAC) framework. (a procedure to do this would simply construct a (hyper-)rectangle around the samples provided: see [31] for further details)*.

## B Additional Details on Experiments

### B.1 Prompts

We distinguish between 2 types of prompts for the API calls to the LLM (in this paper, `GPT`):

- System prompt: We use this prompt to guide the model’s behaviour and responses. It sets the overall instructions for the model, such as defining its role and the syntactic format in which it should respond. In this work, we use this as: “You are a scientist specialising in chemistry and drug design. Your task is to generate valid SMILES strings as a comma-separated list inside square brackets. Return the response as plain text without any formatting, backticks, or explanations. The response must be formatted exactly as follows: [SMILES1, SMILES2, …]. Avoid any extra text or explanations.”
- User prompt: This is the input provided by the user, containing the actual query constructed in manner described below. The LLM generates responses based on this input while considering the instructions set by the system prompt. There are two kinds of user prompts based on whether a set of inhibitors are shown to the LLM during search and generation or not. For revealing known inhibitors, we use: “Generate up to *s* novel valid molecules similar to the following positive molecules: […]”. Otherwise, the prompt is simply “Generate up to *s* novel valid molecules”. We also allow feasible molecules generated in `GenMol` to be used as “context”. In this case, we use a it as a part of the user prompt as: “Additionally, consider these previously generated feasible molecules: […].”

## Footnotes

1 For example, given a sample of feasible molecules, simply find the intervals from the minimum and maximum values of *Affinity* and *SynthesisSteps*.

2 A sample-based estimate is also used in [20] and [21].

3 The most general approach is a branch-and-bound search. However, to be effective for large values of *s*, this will require obtaining an upper-bound on the score obtainable by experiments subsumed by the bounding hyper-rectangle. This is not straightforward for the *Q*-heuristic, unless certain strong assumptions are made.

4 This procedure is inefficient since the generator is invoked for with every element of *E*_*k*_. This can be improved by: (a) Prderomg the elements of *E*_*k*_ by decreasing *Q*-values; and (b) Finding the first element in this sorted sequence for which LMLFStar returns a non-empty set. LMLFStar need not be invoked for subsequent elements in the sequence.

